# Contextual Autocomplete: A Novel User Interface Using Machine Learning to Improve Ontology Usage and Structured Data Capture for Presenting Problems in the Emergency Department

**DOI:** 10.1101/127092

**Authors:** Nathaniel R. Greenbaum, Yacine Jernite, Yoni Halpern, Shelley Calder, Larry A. Nathanson, David Sontag, Steven Horng

**Affiliations:** Beth Israel Deaconess Medical Center — Harvard Medical School — Department of Emergency Medicine, Boston, MA; New York University, New York, NY

**Keywords:** Autocomplete, Contextual Autocomplete, Ontology, Structured Data, Emergency Medicine, Machine Learning

## Abstract

**Objective:** To determine the effect of contextual autocomplete, a user interface that uses machine learning, on the efficiency and quality of documentation of presenting problems (chief complaints) in the emergency department (ED).

**Materials and Methods:** We used contextual autocomplete, a user interface that ranks concepts by their predicted probability, to help nurses enter data about a patient’s reason for visiting the ED. Predicted probabilities were calculated using a previously derived model based on triage vital signs and a brief free text note. We evaluated the percentage and quality of structured data captured using a prospective before-and-after study design.

**Results:** A total of 279,231 patient encounters were analyzed. Structured data capture improved from 26.2% to 97.2% (p<0.0001). During the post-implementation period, presenting problems were more complete (3.35 vs 3.66; p=0.0004), as precise (3.59 vs. 3.74; p=0.1), and higher in overall quality (3.38 vs. 3.72; p=0.0002). Our system reduced the mean number of keystrokes required to document a presenting problem from 11.6 to 0.6 (p<0.0001), a 95% improvement.

**Discussion:** We have demonstrated a technique that captures structured data on nearly all patients. We estimate that our system reduces the number of man-hours required annually to type presenting problems at our institution from 92.5 hours to 4.8 hours.

**Conclusion:** Implementation of a contextual autocomplete system resulted in improved structured data capture, ontology usage compliance, and data quality.

## 1. Introduction

### Background and Significance

Unstructured data accounts for the vast majority of data stored by hospitals. Though plentiful, extracting information and knowledge from this material can be perilous. Structured data is easier to process and analyze, but has been difficult to collect reliably.

The benefits of using structured ontologies is well established. [1] Ontologies form the basis for clinical decision support systems[2], enable research [3], and are essential for interoperability.[4]

However, adoption and use of structured ontologies has been slow. Clinicians often prefer the ease of using natural language [5] since any information system underrepresents the complexity of reality. [6] No classification system can capture the subtlety, nuance, and granularity that exists in the natural world. Redundant entry of patient data, as is often required for ontologies, further stretches limited resources.[7] Browsing existing vocabularies has also proven to be problematic, leading users to select wrong tems. [8] Common solutions, such as tree structure browsers, force users to laboriously read and exclude many irrelevant items to navigate an ontology. Dozens of complex visualization schemes have been proposed, but none have garnered meaningful adoption. [9]

More intelligent, contextual user interfaces can help mitigate these challenges. For example, predictive autocomplete has become a standard feature of most smartphones. The phone displays words or phrases based on the user’s prior conversations and writing style. [10] The software adjusts its suggestions based on context — such as the application that is being used and the person that is being communicating with. In this manner, the phone will suggest more formal terms when writing an email to a colleague, and adopt a more casual tone when sending a text message to a friend.

Similarly, in our autocomplete implementation, we used contextual patient information to predict a patient’s presenting problem (chief complaint). Our predictions were based on the patient’s vital signs and the nurses’ description of their state (e.g. medical history, symptoms) at arrival.[11] The more that is known about the patient, the more accurate our autocomplete suggestions.

Prior work by Sevenster et al.[12,13] modeled the utility of various autocomplete algorithms to search SNOMED terms. Their approach worked best when MEDLINE abstracts were used to compute the semantic distance between terms and then a multi-word matching technique was applied to obtain a theoretical 7% to 18% reduction in keystrokes. Our approach differs in that we deployed autocomplete based on contextual patient information in a live clinical setting.

In our study, we used emergency department (ED) presenting problem data as an example for how structured data capture can be improved with contextual autocomplete. The presenting problem is a critical component of emergency medicine decision making, setting the stage and priorities of the ED visit. When patients present to the ED, they share a reason for their visit with the initial provider. These first spoken words are the chief complaint, which is then interpreted by the triage nurse and recorded as the presenting problem. Traditionally, this data has been unstructured free text — filled with variations in wording, spelling, and descriptiveness across different sites and providers.

The ubiquity and heterogeneity of unstructured data is a major hindrance to automating and improving healthcare. Contextual autcomplete can be applied more broadly to other unstructured data problems in healthcare, as there are very few areas where there is no prior distribution on terms. Notably, we show that structured data collection can be made easier than unstructured data collection techniques. Such a system encourages ontology usage, enhances secondary data use, enables clinical decision support, and automates workflows. [14]

### Objective

In this study we sought to determine the effect of contextual autocomplete, a user interface that users machine learning, on the efficiency and quality of documentation of presenting problems in the emergency department. We did this by measuring percentage of structured data capture, completeness, quality, and precision of the data, as well as number of keystrokes required for data entry.

## 2. MATERIALS AND METHODS

### Study Design

We performed a mixed methods study that consisted of both a prospective cohort before-and-after design and a qualitative study.

Our study was comprised of a 12 month pre-intervention period, a 16 month postimplementation phase, and a 32 month development period between while our contextual autocomplete system was being developed.

During the post-implementation phase, our system experienced several periods of unplanned downtime. We performed a subgroup analysis to compare the percentage of presenting problems properly mapped to our ontology while our prediction system was on and while it was off. No other variables were altered during this period. When the prediction system is not available, the autocomplete falls back to ordering terms by alphabetical order. Therefore, this is subgroup analysis is a comparison between contextual autocomplete and standard autocomplete.

### Setting and Selection of Participants

We prospectively captured data on 279,231 consecutive emergency department patient encounters at an urban, academic, Level I trauma center from September 2011 to September 2016. No patients were excluded. (TABLE 1)

**Table 1:** Demographics

|  | Pre-Implementation |  | Development |  | Post-Implementation |  |
| --- | --- | --- | --- | --- | --- | --- |
| Approx. Dates | Sept. 2011 - Sept. 2012 |  | Sept. 2012 - May 2015 |  | May 2015 - Sept. 2016 |  |
|  |  |  |  |  | Autocomplete Off | Autocomplete On |
| n (%) | 56,186 (20.2%) |  | 144,077 (51.8%) |  | 854 (0.3%) | 77,303 (27.8%) |
| Mean Age | 55.1 |  | 54.2 |  | 50.3 | 52.7 |
| Sex (% female) | 54.8% |  | 54.5% |  | 52.6% | 54.4% |
| **Acuity<sup>†</sup>** | 3 [2-3] |  | 3 [2-3] |  | 3 [2-3] | 3 [2-3] |

**†Median Emergency Severity Index (ESI) with interquartile ranges.**

### Predictive Model

We previously developed an algorithm [11] to suggest presenting problems from a controlled vocabulary based on triage vital signs and the brief triage note entered by the nurse when the patient first presents to the emergency department. This model was developed using 97,000 triage notes using a multiclass Support Vector Machine (SVM), representing free text as a bag-of-words. The model had a best-5 accuracy of 0.757, which represents how often the list of most likely predicted labels actually contained all of the true presenting problems.

### Intervention

Our contextual autocomplete system ranked concepts by their predicted probability, not alphabetical order. By displaying only the most relevant concepts, we eliminated much of the searching associated with traditional ontology user interfaces.

We favored a ‘contains search’ over a ‘begins with search’. In a ‘begins with search’ only the leading characters are used to match terms; a ‘contains search’ will match the characters searched for if they appear anywhere in the term. For example, if nurse searches for *‘DVT’*, a medical abbreviation for deep venous thrombosis, a ‘begins with search’ will not locate the concept ‘Leg *DVT’* owing to the fact that the concept does not begin with the characters *‘DVT’*. Conversely, a contains search for *‘chest pain’* will successfully locate ‘pleuritic *chest pain’*, a subtype of chest pain.

The five ‘best guesses’ generated by the system were also presented to the user. The user was then able to click on one or more presenting problems from the list created by our system. If the user did not see their desired presenting problem in the list, they could begin typing the first few letters of the presenting problem they desired and our system would refine the top five best guesses based on the user’s input. The user was also able to ignore our system’s recommendations and enter their own free text presenting problems. (Figure 1)

**Figure 1:**
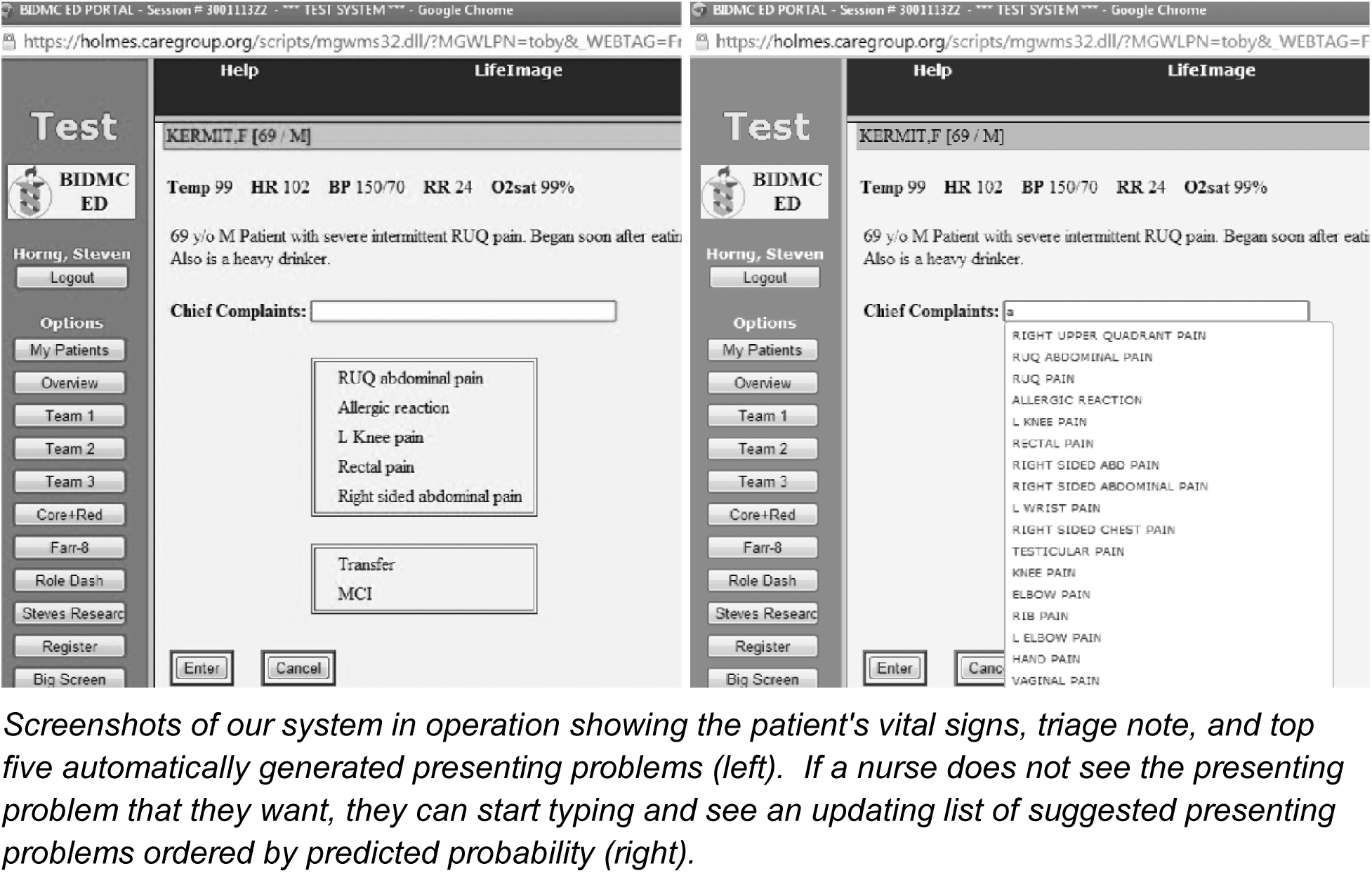
User Interface Screenshots

### Data Collection and Processing

We divided our data into three categories representing the time period prior to our intervention, the time period during our system’s development, and the time period post-implementation. The post-implementation period was further analyzed based on whether our system was operational or not on a per-patient basis.

We also conducted a qualitative review of 150 patients. We randomly sampled ten patients from each of the five Emergency Severity Index (ESI) levels during both the pre-intervention and and post-intervention periods; this yielded a total of 100 records representing a broad spectrum of presenting problems. To this pool we added 50 randomly selected charts that had been completed during our unscheduled downtime.

Three independent, expert reviewers qualitatively reviewed each of the 150 patient’s records. Reviewers were blinded to the selection criteria for the patients and the results of our primary outcome measure.

### Outcome Measures

Assessment of our primary outcome measure occurred at the level of the patient. Since each patient may have more than one presenting problem, we defined the primary outcome measure as positive if *all* of the documented presenting problems listed for the patient were able to be mapped to our ontology. If any of the presenting problems were not mapped (i.e., the triage nurse used free text or a term not in our ontology), we considered the outcome to be negative. For example, the entire patient visit with a presenting problem of “nausea, other” would be considered negative because “other” is not in the ontology. A patient visit of “nausea,vomiting” would be considered positive because all concepts are in the ontology. This all or none strategy provides the most conservative estimate of our system’s performance.

For the qualitative component, all reviewers independently assessed every chart’s presenting problem for completeness, precision, and overall quality. Each of these metrics was scored on a four-point likert scale.

### Primary Data Analysis

For each time period we computed the percentage of patient encounter in which all of the patient’s presenting problems were successfully mapped to our ontology. For the qualitative component, reviewer’s scores were aggregated and means computed for each data element outlined above.

We also conducted a subgroup analysis of our system’s performance during unscheduled downtime when autocomplete was turned off, measured by the lack of recorded predicted presenting problems in the system. Since predictions can also be missing when patients skip triage, we excluded patients that did not have a triage note (1.0%). (FIGURE 2) These typically represent the sickest patients such as those in cardiac arrest, suspected strokes, and trauma. Due to the severity of their disease, these patients often bypass triage. We also conducted a sensitivity analysis in which we placed all 811 excluded patients back in, which did not show a clinically or statistically significant difference.

Lastly, we calculated the number of keystrokes used to type each presenting problem during each of the study periods. Given the chaotic environment of Emergency Department triage, measuring absolute time would not be representative as triage nurses often perform several tasks simultaneously and are interrupted frequently. It is therefore difficult to reliably attribute duration to any one single activity. We chose instead the surrogate measure of keystrokes entered or ‘number needed to type’ for the amount of time that was spent documenting presenting problems.

**Figure 2:**
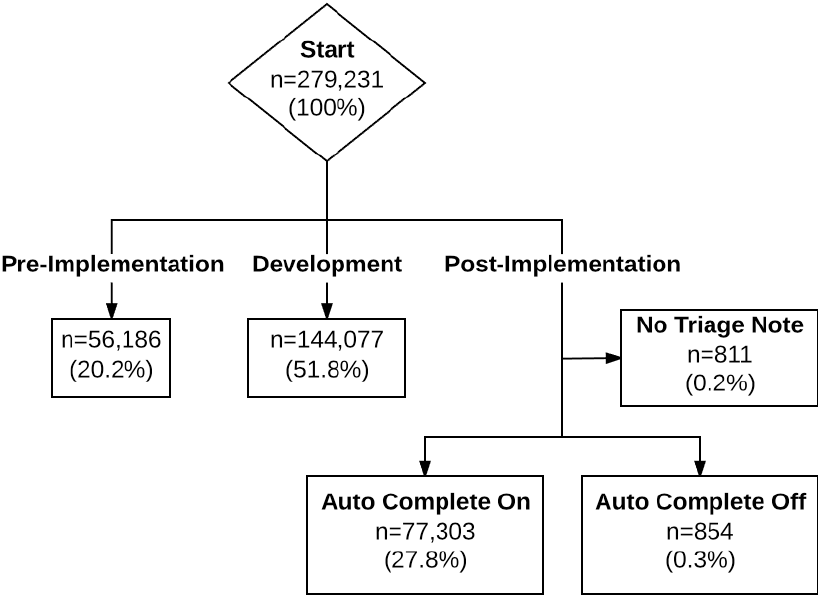
Enrollment Diagram

## 3. RESULTS

Prior to implementing our system 26.2% of presenting problems entered at triage were able to be automatically mapped to a structured ontology. After deployment, while our system was operational, we improved performance to 97.2% (p<0.0001). (TABLE 2; FIGURE 3)

**Table 2:**
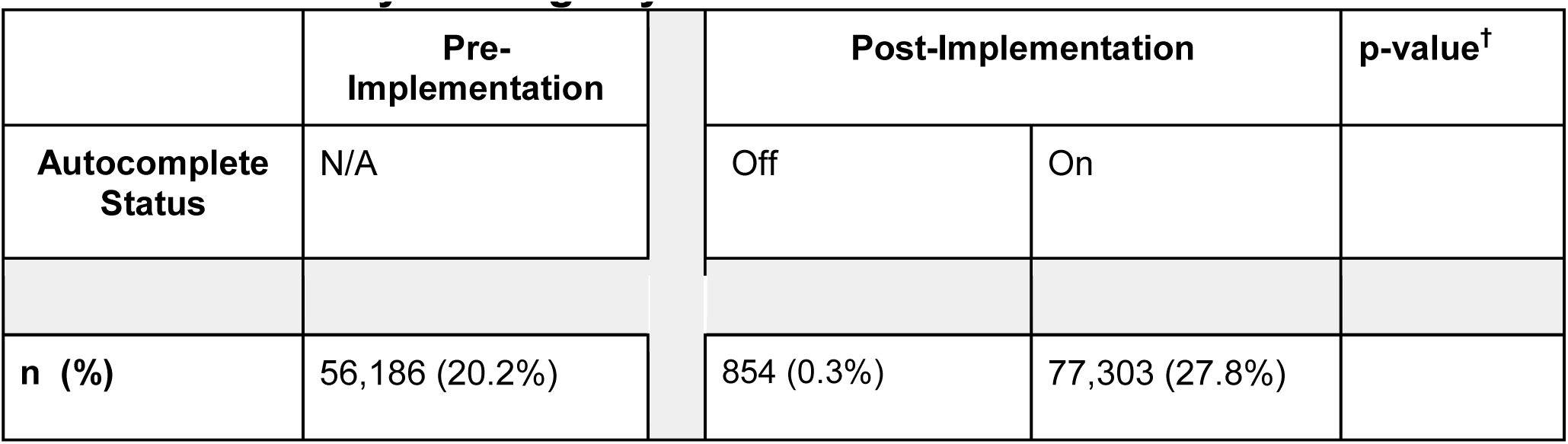
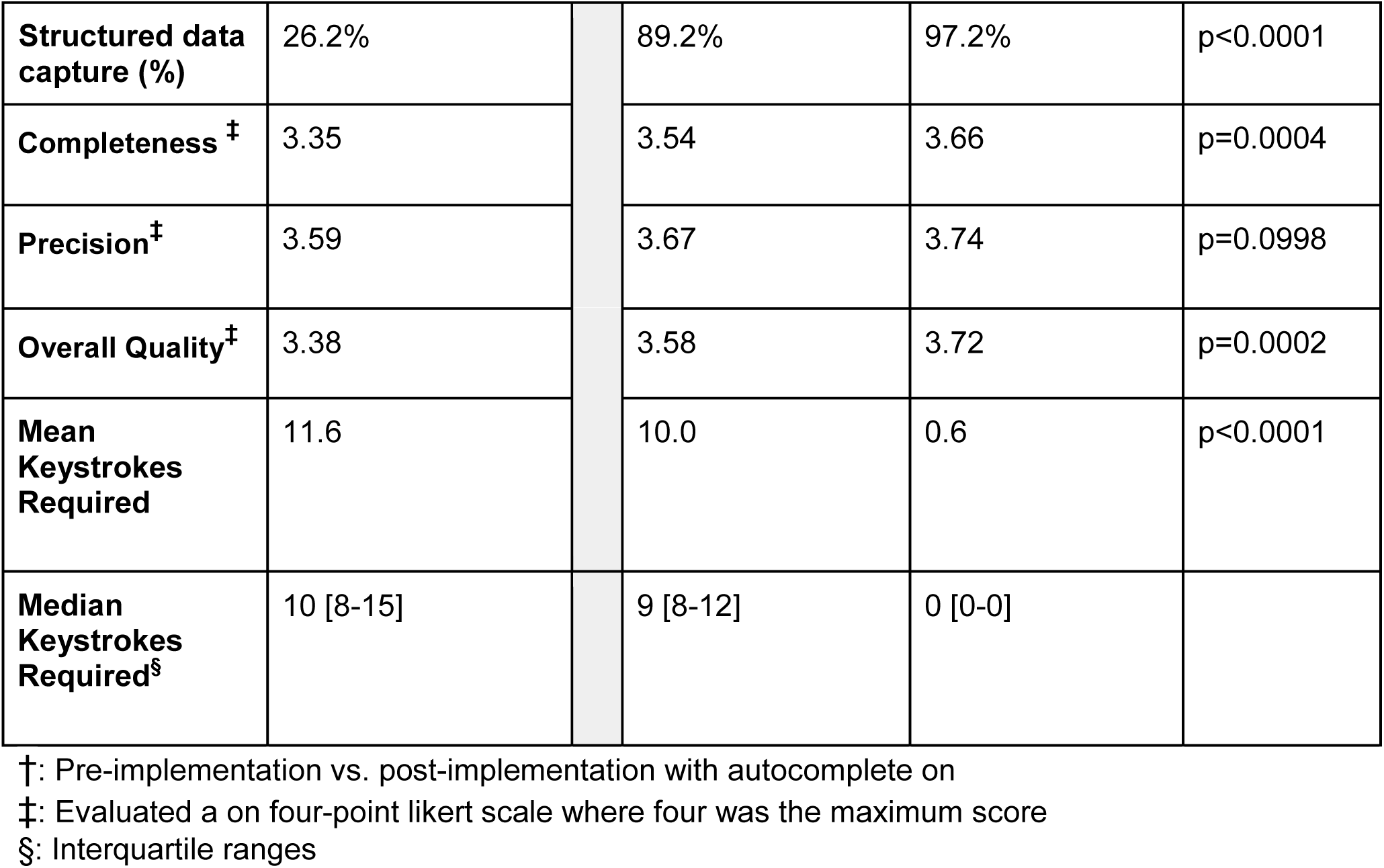
Summary Findings by Time Period

**FIGURE 3:**
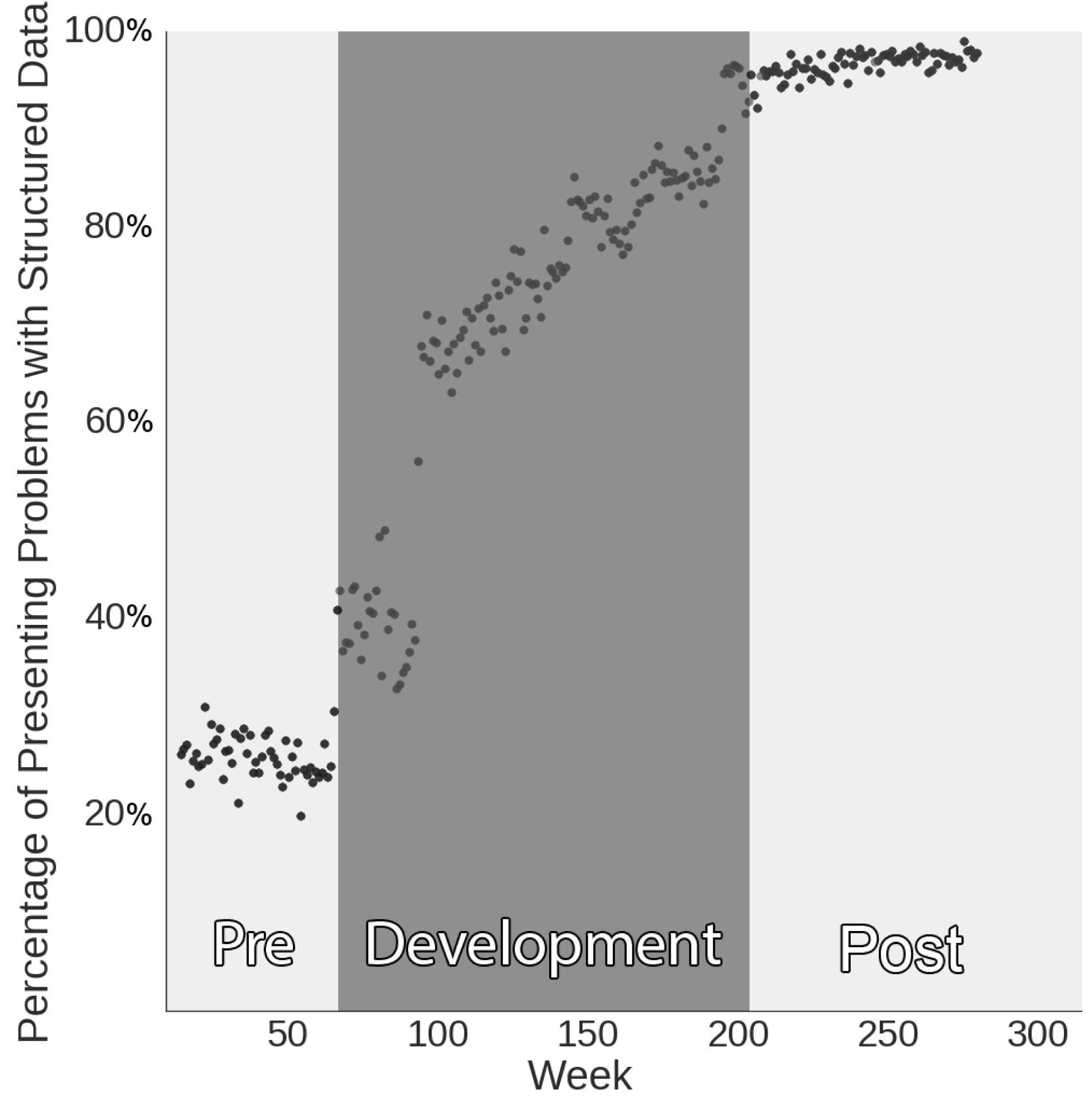
Percentage of Presenting Problems Mapped to Structured Data by Week

During the post-implementation phase, while the system was operational, an average of 1.48 presenting problems were documented for each patient. To record these, nurses clicked the auto-suggested presenting problem 88.7% of the time. Users erroneously clicked, and subsequently deleted, 3.8% of complaints.

Qualitative assessment of presenting problem showed contextual autocomplete was associated with more complete (3.35 vs 3.66; p<0.05), equally precise (3.59 vs. 3.74; p=0.1), and higher overall quality (3.38 vs. 3.72; p<0.05) presenting problems.

Our system experienced multiple episodes of unscheduled downtime, which are not correlated with patients or staff. We aggregated these incidents of downtime to give us an estimate of how triage nurses perform with non-predictive autocomplete alone. During these episodes 854 patients were triaged; only 89.2% of these encounters resulted in structured data capture.

During our pre-implementation phase the estimated mean number of keystrokes typed for each presenting problem was 11.6. In the post-implementation phase with our system active we reduced the estimated mean ‘Number Needed to Type’ to 0.6 characters, a 95% improvement. (FIGURE 4)

**FIGURE 4:**
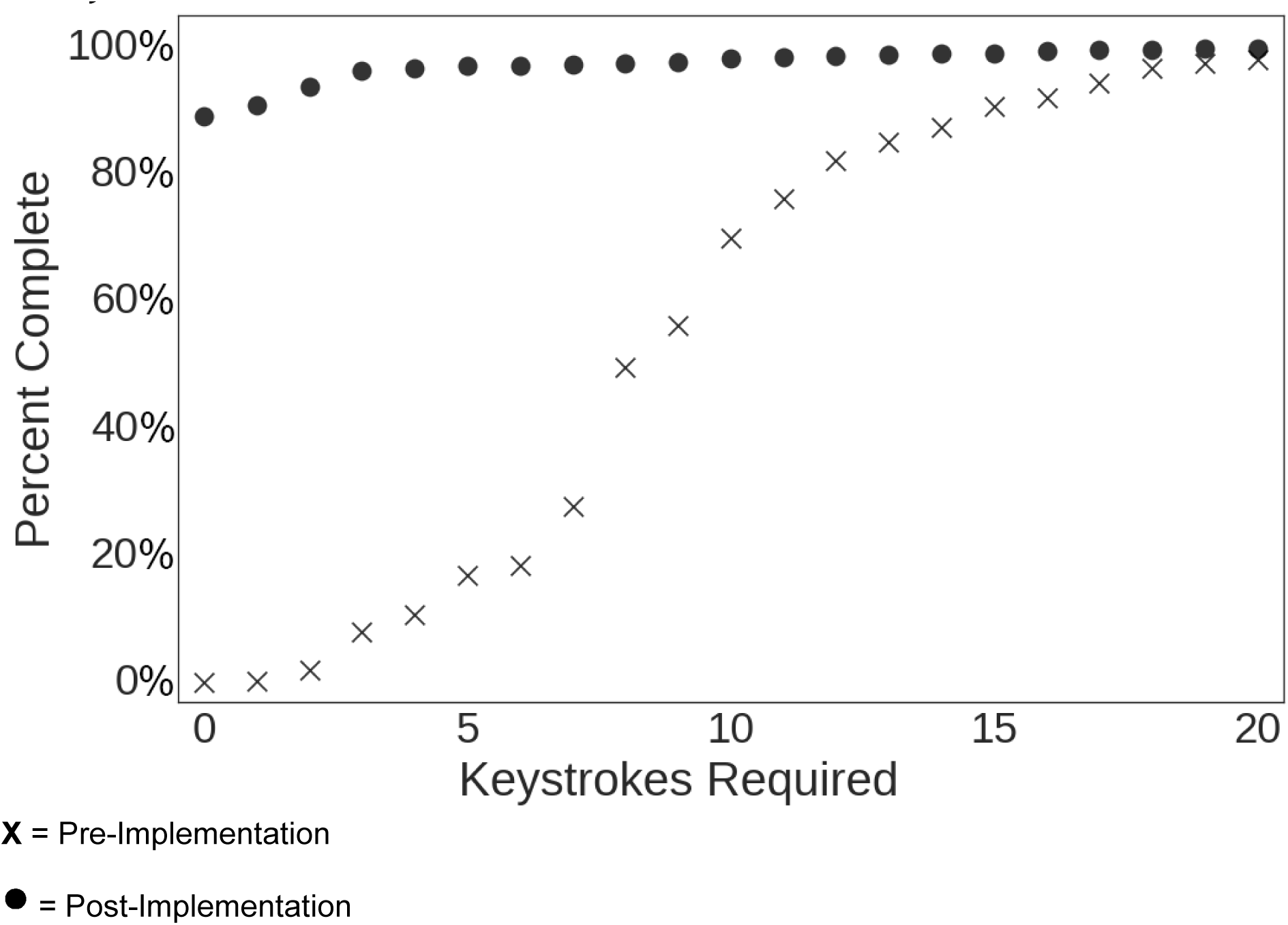
Number of Keystrokes Required to Complete Data Entry

## 4. DISCUSSION

As part of a multi-year quality improvement project, our emergency department sought to standardize nursing presenting problems to enable presenting problem based clinical workflows and retrospective analysis. Hundreds of hours were spent by nursing administration on training, education, and ultimately remediation for nurses non-compliant with the standardized list. Despite the great efforts to standardize the collection of presenting problems, the minimal improvement was transient and unsustainable without constant feedback.

Structured ontology adoption has been impeded by poor user interface design and the time burden required for their use. By deploying a contextual autocomplete system, we have demonstrated a technique that captures structured data on nearly all patients. Our system attained a 97.2% structured data capture rate while simultaneously improving documentation quality and reducing the burden on triage nurses, saving almost 88 hours per year in data entry.

Traditionally, unstructured data has been easy to collect, but extracting knowledge from this source has been arduous. Structured data, which has been harder to obtain, provides appreciable benefits. Using contextual autocomplete, we reversed this paradigm, showing how structured data collection can be made easier and faster than unstructured collection.

Prior work has focused primarily on using natural language processing and machine learning to extract structured data from unstructured text. [15], [16] However, retrospective classification of presenting problems makes assumptions about clinical meaning when there is not an exact match to a term in the ontology. Simply, it is impossible to know if the term the triage nurse would have used is the same as the one predicted retrospectively. In contrast, we present our users with our best guesses of presenting problem and then collect structured data prospectively, ensuring that the selection from the ontology accurately reflects clinical thinking.

Contextual autocomplete reduces the number of keystrokes needed to document a presenting problem to zero for the vast majority of patients, 89.3%. Overall, we reduced the number needed to type by 95%. Given median typing speeds [17] and mean english word length [18], we estimate that our system reduces the number of man-hours required annually to type presenting problems at our institution from 92.5 hours to 4.8 hours. The reduced workload, speed, and ease of use our system provides mitigates many of the challenges that have historically impaired structured ontology use.

In the post-implementation period, we performed a subgroup analysis comparing when the system was on versus off. The percentage of structured data capture decreased from 97.2% on to 89.2% off. On interviews with users, it was noted that there was a strong learning effect over time, where users learned terms in the ontology over time, lessening the transient loss of contextual autocomplete functionality. Given the high staff turnover of users in the emergency department, this learning effect would only be temporary.

## 5. LIMITATIONS

The data utilized in this study was obtained from a single tertiary academic medical center. As a result, our outcomes may not be generalizable to other emergency departments practicing in disparate geographical areas or with a different patient population. We also used an ontology that was developed at the same site as our study. As result, our successful match rate is likely to be higher than would be expected at a different clinical site with a different patient distribution.

## 6. CONCLUSION

We implemented a contextual autocomplete system that exhibited a marked improvement in structured data acquisition and ontology usage compliance. Despite the challenges commonly associated with ontology use, we demonstrate a system that recorded structured data for 97.2% of patients, improved the quality of documentation, and reduced that amount of time required for data entry.

Contextual autocomplete has demonstrated its effectiveness for presenting problems and can almost certainly be expanded to additional areas such as diagnosis, procedures, and problem lists to streamline user data entry and improve ontology adherence.

## 9. ACKNOWLEDGEMENTS

We would like to acknowledge Scott Rollins for his help evaluating the quality of presenting problem documentation.

## 10. CONTRIBUTORS

SH and DAS conceived and designed the study. YJ, YH, and DAS developed the algorithm. SH, SC, and LAN collected the data. SH and NRG performed the analysis. NRG, SH, and DAS drafted the manuscript. All authors contributed substantially to its revision. SH takes responsibility for the paper as a whole.

